# Impact of Meteorological factors on wheat growth period and irrigation water requirement —A case study of the Beijing-Tianjin-Hebei region in China

**DOI:** 10.1101/2021.01.04.425202

**Authors:** Chengcheng Xu, Chuiyu Lu, Jianhua Wang

## Abstract

This study analyzes the irrigation water requirement of wheat in the growth stage in the Beijing-Tianjin-Hebei region under the changes of meteorological factors conditions, using the growth period data, and meteorological data from 80 meteorological stations, from 2000 to 2019. The results show that: (1) The annual average precipitation, average wind speed, and average relative humidity of the growth period in the Beijing-Tianjin-Hebei region show a downward variation trend, while the temperature variation shows an upward trend. Moreover, relative humidity and radiation exhibit a negative spatial correlation. (2) Wheat irrigation water requirement in the Beijing-Tianjin-Hebei region gradually decreases from north to south and east to west. However, the eastern region shows a gradually increasing trend, while the western region shows a decreasing trend. (3) Meteorological factors are negatively correlated with irrigation water requirement, potential evapotranspiration, effective precipitation, and relative humidity, and significantly positively correlated with sunshine hours, average temperature, and wind speed. The overall variation in irrigation water requirement has the highest correlation with potential evapotranspiration. However, the yearly variations in regional irrigation water requirement are dependent on factors such as wind speed, relative humidity, and radiation.

## 1. Introducion

The United Nations defines climate change as the variation in the Earth’s atmospheric composition from direct or indirect human activities, in addition to the natural variability of climate observed during similar periods. The impact of climate change on the human environment has increased progressively and extensively (Seo et al., 2013; Ju et al., 2013; Calzadilla et al., 2013; Noufé et al., 2015), and since the First World Climate Conference in 1979, climate change research has gradually attracted global attention (Iglesias, 2012; Ahmad et al., 2019; Mayank et al., 2019; Pons et al., 2020). The World Meteorological Organization (WMO) and the United Nations Environment Programme (UNEP) are two major organizations dealing with climate change, that simultaneously established the Intergovernmental Panel on Climate Change (IPCC) (Van der and Warner, 2020). The various assessment reports put forward by the IPCC since 1990 have established a scientific basis for climate change research, by indicating that global warming was caused by both natural and manmade factors. The global temperature reportedly rose by 0.3-0.6 °C in a century. Furthermore, the elevation in global average temperature since the mid-20th century was primarily attributed to greenhouse gases emitted through human activities. The global climate change since the past 132 years (1880-2012) was also elaborated. Moreover, the IPCC 2013 report predicts the trend of climate change at the end of the 21st century through a variety of scenario models: Temperature rise varies regionally, and it will be higher on the land than the sea by 1.4-1.7 °C, with the Arctic recording the highest temperature (McBride et al., 2020). Extreme hot weather events will increase, and the duration will be lengthened in most areas, and extreme cold weather events will decrease. The frequency of high temperature events will double every two decades. Increased precipitation will occur mostly in high latitudes and some in mid latitudes, instead of the subtropics. Precipitation will increase with the rise in global warming, and the rate of increase in precipitation for a unit temperature rise is less than that of water vapor. Precipitation will show high spatial variations, further increasing the precipitation gap between arid and humid regions (Weerathunga et al., 2020), and in most parts of the world, between wet and dry seasons. In addition, precipitation will increase in the equatorial Pacific, high latitude and humid mid-latitude regions and decrease in subtropical and arid mid-latitude regions.

The agriculture industry is most vulnerable to the effects of climate change (Sun et al., 2013; Schönhart et al., 2016; Liu et al., 2016). Therefore, the impact of climate change on agricultural production is a relevant issue in climate change research (Dong et al., 2019). Climate change in China follows a similar trend as the overall global climate change, with slight variations (Lv et al., 2013; Zhang et al., 2016). The North China plain is one of the principal wheat producing areas in China. Climate change inevitably impacts the growth period and water requirement of wheat, thereby affecting its yield and quality (Chen et al., 2013; Wang et al., 2014; Geng et al., 2019; Wang et al., 2019). Thus, the study of regional changes in irrigation water requirement for wheat in the Beijing-Tianjin-Hebei region is significant in relation to the overall temperature rise in North China. Furthermore, the factors affecting wheat irrigation water requirement have also been investigated.

## 2. Materials and methods

### 2.1 Overview of the study area

The Beijing-Tianjin-Hebei region (total area of 216,500 km^2^), is the “capital economic circle” of China and accounts for 2.3% of the total area of the country. In addition to Beijing and Tianjin, the region comprises Baoding, Langfang, Shijiazhuang, Tangshan, Handan, Qinhuangdao, 11 prefecture-level cities including Zhangjiakou, Chengde, Cangzhou, Xingtai, and Hengshui, and 2 provincial-level cities directly under the control of Dingzhou and Xinji, of the Hebei province. The Taihang Mountains lie to the west, and the Bohai Bay to the east of the Beijing-Tianjin-Hebei region. The terrain is high in the northwest and north, and flat in the south and east (Zhang et al., 2020). In winter, the cold and snowless weather is controlled by the Siberian continental air mass, and northerly and northwestern winds prevail. Spring is affected by the Mongolian continental air mass, such that there is a rapid temperature rise, high wind speed (Guo et al., 2020), dry climate, and amount of evaporation. The weather tends to be dry, windy, and sandy (Guo et al., 2020). Summer is affected by oceanic air masses, and is humid, with high temperatures, and heavy rainfall (Song et al., 2020). However, the inconsistencies in time, intensity, and range of influence of the Pacific subtropical high every summer result in high rainfall variation and occasional droughts and floods. In autumn, the weather is high and the rainfall is low (Zhenyu et al., 2019; Men et al., 2020; Bi et al., 2020).

In the Beijing-Tianjin-Hebei region, 13 main single crops are cultivated, including wheat, corn, cotton, vegetables, potatoes, peanuts, millet, soybeans, forest fruits, and rice. Multiple cropping occupies a part of the region, and is observed primarily for wheat and corn, in addition to wheat and peanuts, soybeans and cotton, cotton and vegetables, and cotton and fruits. Other types of crops such as rape, flax, naked oats, and sorghum, are also cultivated in small areas. However, wheat and corn are the primary plantations occupying a large portion of the Beijing-Tianjin-Hebei region.

### 2.2 Source of information

The meteorological data were obtained from the National Meteorological Center of China. Daily meteorological data, including average temperature (°C), maximum temperature (°C), minimum temperature (°C), sunshine hours (h), precipitation (mm), wind speed (m/s), and average relative humidity (%) were selected from 80 representative ground weather stations from 2000 to 2019. For unavailable meteorological data, the missing values of temperature (average, maximum, and minimum) and precipitation were interpolated using Kriging interpolation.

The growth period data for wheat was obtained from the Chengde, Tangshan, Zunhua, Baoding, Shijiazhuang, and Botou observation stations from 2000 to 2019. Since the growth period observation stations and ground weather stations (Beijing Station, Tianjin Station, Shijiazhuang Station, and Nangong Station) were not consistent, the nearest matching principle was adopted and 80 sets of continuous 20-year observation data were selected, as shown in Table 1.

**Table 1.**
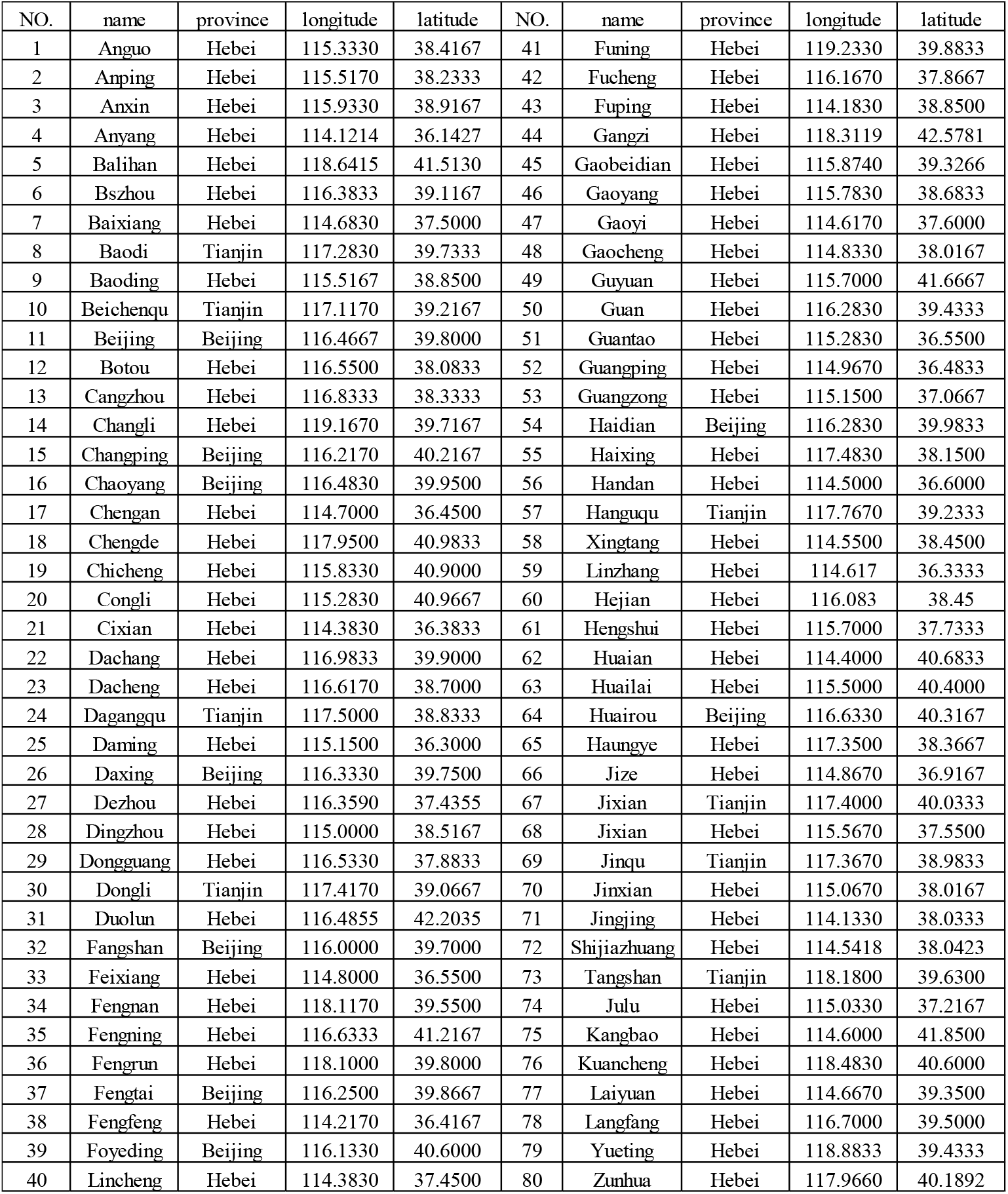
Data of Beijing-Tianjin-Hebei meteorological stations

This study divided the entire growth period of wheat into six stages: initial growth period, thawing period, wintering period, fast development period, middle growth period, and maturity period. The multi-year average of the development period was used as the local average growth period.

### 2.3 Calculation of potential evapotranspiration

Potential evapotranspiration (ET_0_) is the maximum evapotranspiration that a fixed underlying surface can reach when the water supply is not restricted by meteorological factors (Lecina et al., 2003). It determines the dry and wet conditions of the region, along with precipitation, and is a key factor for estimating the ecological water requirement and agricultural irrigation. Further, in the study, the characteristics of the crop are specified, by defining the evapotranspiration of a green grass canopy with the same height, moderate moisture, active growth, and complete coverage of the ground surface with a crop height of 0.12 m, a leaf resistance of 70 s/m, and a reflectivity of 0.23.

ET_0_ is based on the daily values of average, maximum and, minimum temperatures, average wind speed, average humidity and sunshine hours. It is calculated using the Penman-Monteith method recommended by FAO (Allen et al., 1998):

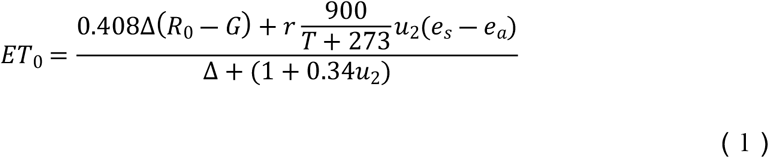

In the equation, *R*_0_ is the net radiation on the surface of the canopy (MJ/(m^2^·d)); *G* is the soil heat flux (MJ/(m^2^·d)); *T* is the average temperature (°C); *e_s_* is the saturated vapor pressure (kPa); *e_a_* is the actual vapor pressure (kPa); Δ is the tangent slope of the saturated vapor pressure and temperature curve at T (kPa/°C); *r* is the hygrometer constant (kPa/°C); *u*_2_ is the wind speed at a height of 2 m (m/s).

### 2.4 Calculation of effective precipitation

Effective precipitation refers to the precipitation required to balance crop evapotranspiration during the growth period. It is related to the crop water requirement and yearly distribution of precipitation. There are differences in the amount of precipitation that specific crops can use for the same annual precipitation and distribution of different precipitation years, as shown in Figure 2. The crop water requirement that is not met by effective precipitation needs to be supplemented through irrigation, Hence, there is a correlation between effective precipitation and irrigation water requirement.

**Figure 1.**
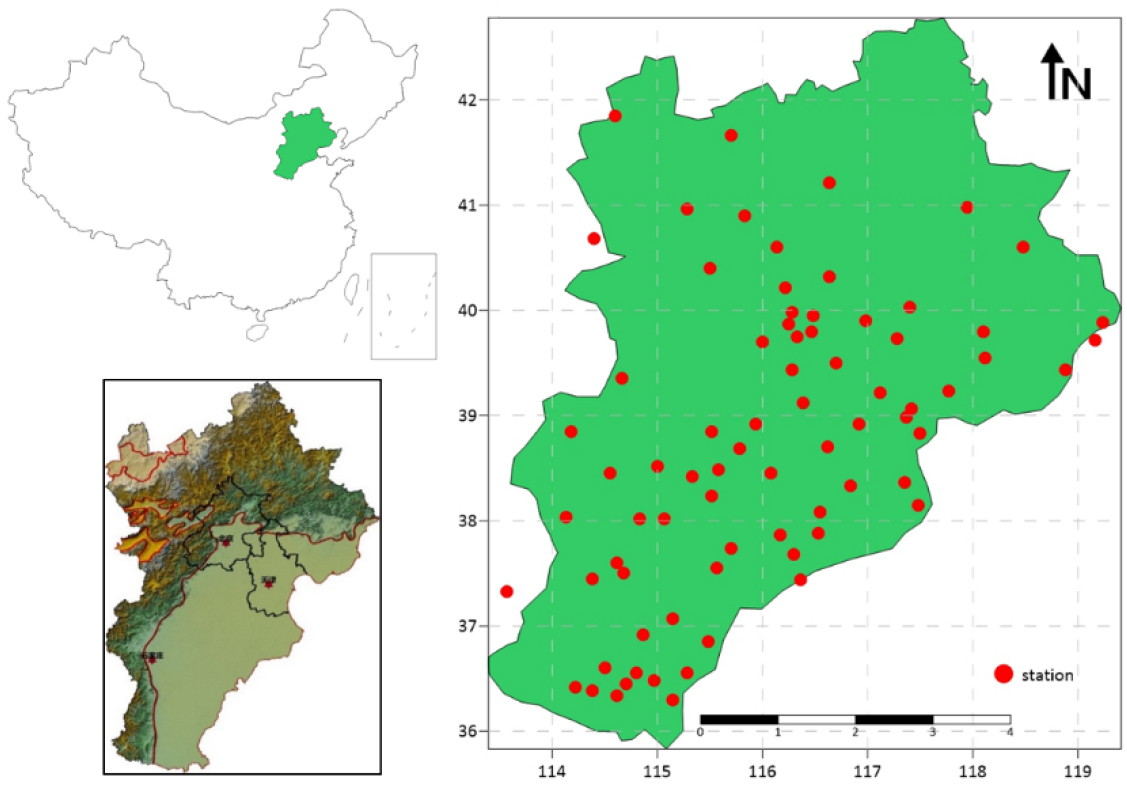
(a) Map showing geographical location of the study area and (b) Map showing distribution of meteorological stations.

**Figure 2.**
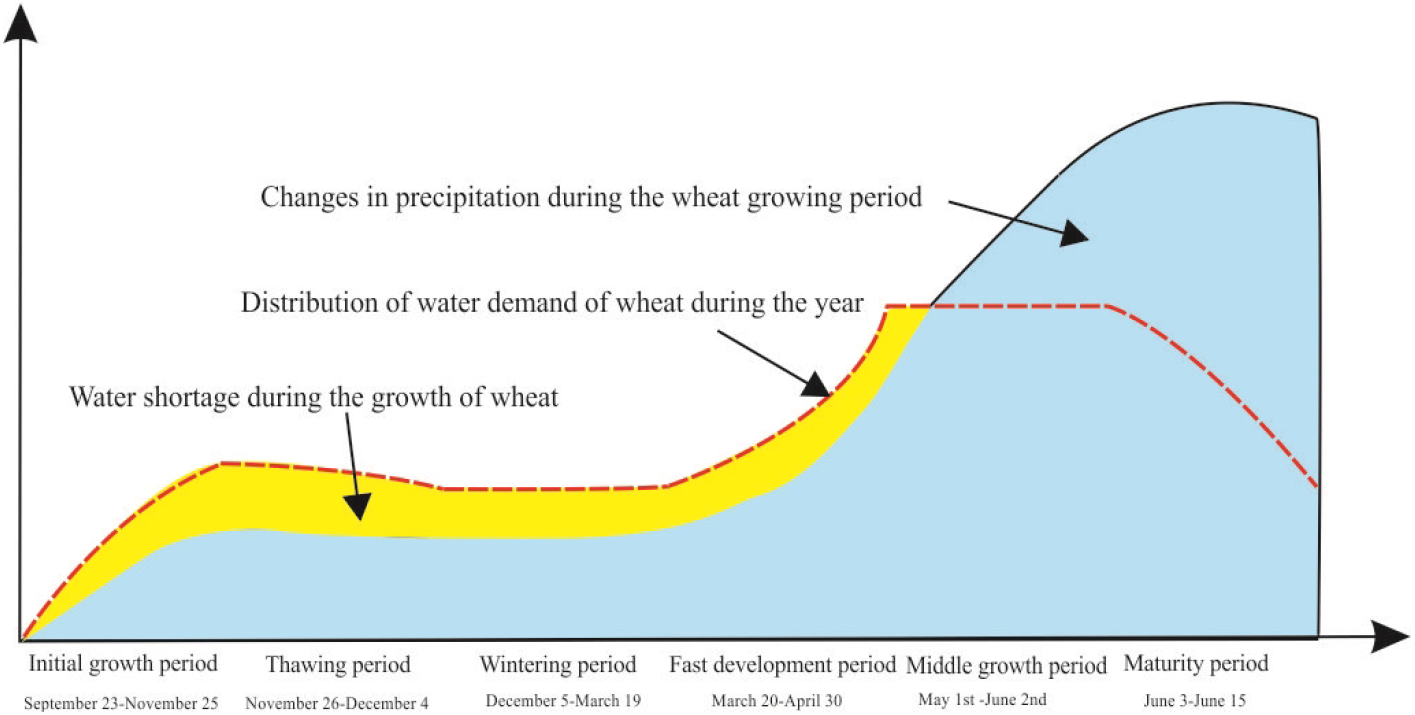
Plot showing the relationship between water requirement and precipitation during wheat growth period.

The crop water requirement is determined by the crop coefficient method as follows:

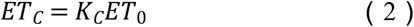

In the formula, *ET_C_* is the crop water requirement (mm/d); *ET*_0_ is the reference crop evapotranspiration (mm/d); and *K_C_* is the crop coefficient.

The piecewise single-value average crop coefficient method recommended by the Food and Agriculture Organization (FAO), is adopted for *K_C_*. Hence, the variation of *K_C_* during the entire growth period is divided into 6 stages. The values of *K_C_* are shown in Table 2.

**Table 2.** Wheat crop coefficients for the study area

| Wheat | Initial growth period | Thawing period | Wintering period | Fast development period | Middle growth period | Maturity period |
| --- | --- | --- | --- | --- | --- | --- |
| Date | September 23-November 25 | November 26-December 4 | December 5-March 19 | March 20-April 30 | May 1-June 2 | June 3-June 15 |
| $K_c$ | 0.6 | 0.6-0.4 | 0.4 | 0.4-1.15 | 1.15 | 1.15-0.4 |

High precipitation occurring in a day may be stored in the soil and subsequently utilized by crops. Thus, effective precipitation can be statistically characterized over time. In this study, the water requirement and characteristics of crop growth every ten days are used for the statistical calculation of effective precipitation as:

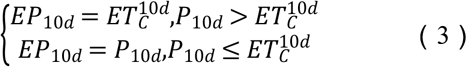

where, *EP*_10*d*_ is the ten-day scale effective precipitation (mm/10 d); 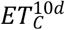 is the ten-day crop water requirement (mm/10 d); and*P*_10*d*_ is the ten-day precipitation (mm/10 d).

The effective precipitation during the crop growth cycle is the sum of the effective precipitation on the ten-day scale:

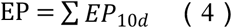

### 2.5 Calculation of irrigation water requirement

A portion of the water needed for agricultural production is obtained from precipitation, and the remaining portion from artificial irrigation. Precipitation is sufficient for crop growth during high-water period, but artificial irrigation is required during low-water period. Therefore, irrigation water requirement, effective precipitation during the crop growth and development stages, the actual area, and the amount of precipitation are closely related factors. Setting the effective precipitation as P_1_ and the actual regional precipitation as P_2_, the irrigation water requirement, Q can be expressed as

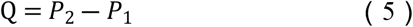

## 3. Result analysis

### 3.1 Analysis of climate change characteristics

#### 3.1.1 Rainfall and temperature

The overall precipitation in the Beijing-Tianjin-Hebei region has exhibited a decreasing trend for over two decades, as shown in Figure 3. There is a negative correlation between precipitation and temperature, the correlation coefficient is −0.65. When the precipitation peaks, the temperature shows a dip.

**Figure 3.**
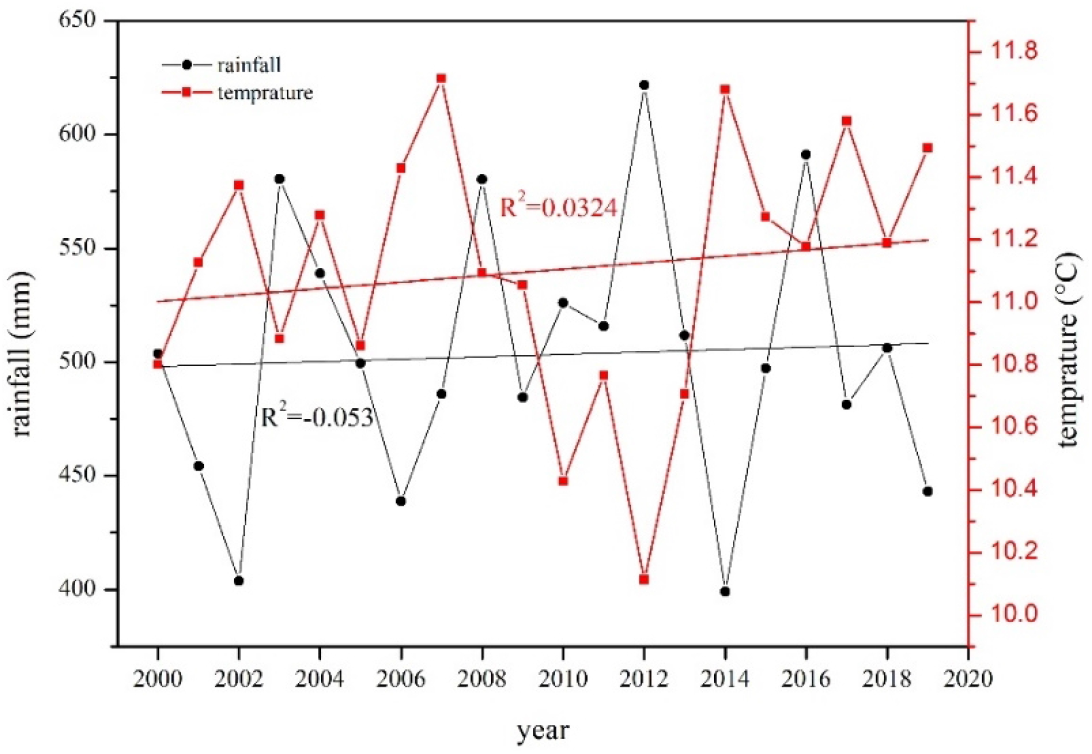
Plot showing the interannual variation trend of rainfall-temperature.

#### 3.1.2 Wind speed, radiation, and relative humidity

From the analysis of the multi-year change from the past 20 years in the Beijing-Tianjin-Hebei region, the wind speed has shown a decreasing trend. From 2000 to the present, the multi-year average wind speed was approximately 2.18 m/s. In 2008, the minimum value was 2.05 m/s. The overall range of radiation showed smooth variations, with increasing and decreasing trends in some years; the overall relative humidity showed a downward trend, with a relative humidity change trend (R^2^) of 0.359, as shown in Figure 4(c). Analyzed from the spatial change trend, the wind speed was higher in the northern part of the study area, and lower in the transition area between the plain and the mountainous regions. When it reached the southeastern plain, the wind speed gradually increased in the coastal zone, as shown in Figure 5(a), and the humidity gradually increased from the northwest to the southeast. The relative humidity in the hilly area was lower than that of the plain area, and high in the southernmost part of the study area, as shown in Figure 5(b). The average radiation hours were higher in the northwest mountainous area. The spatial radiation variation was high in the Southeast Plain, and relatively low in the southernmost part of the study area, as shown in Figure 5(c). Hence, it can be concluded that relative humidity and radiation are inversely correlated in space; that is, areas with high relative humidity have low radiation, and areas with low relative humidity have high radiation.

**Figure 4.**
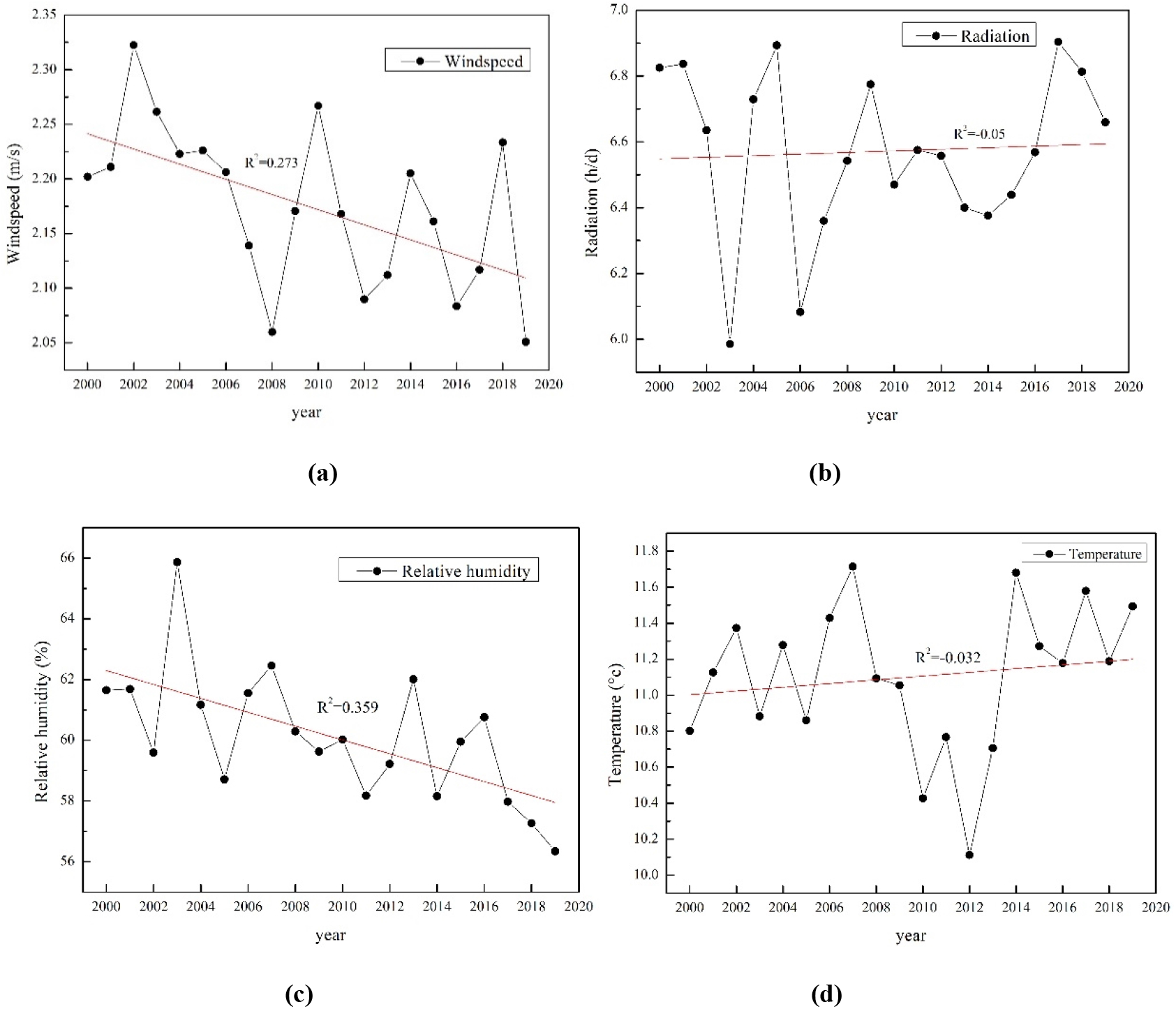
Plot showing the interannual variation trends of (a) wind speed, (b) radiation, (c) relative humidity, and (d) temperature.

**Figure 5.**
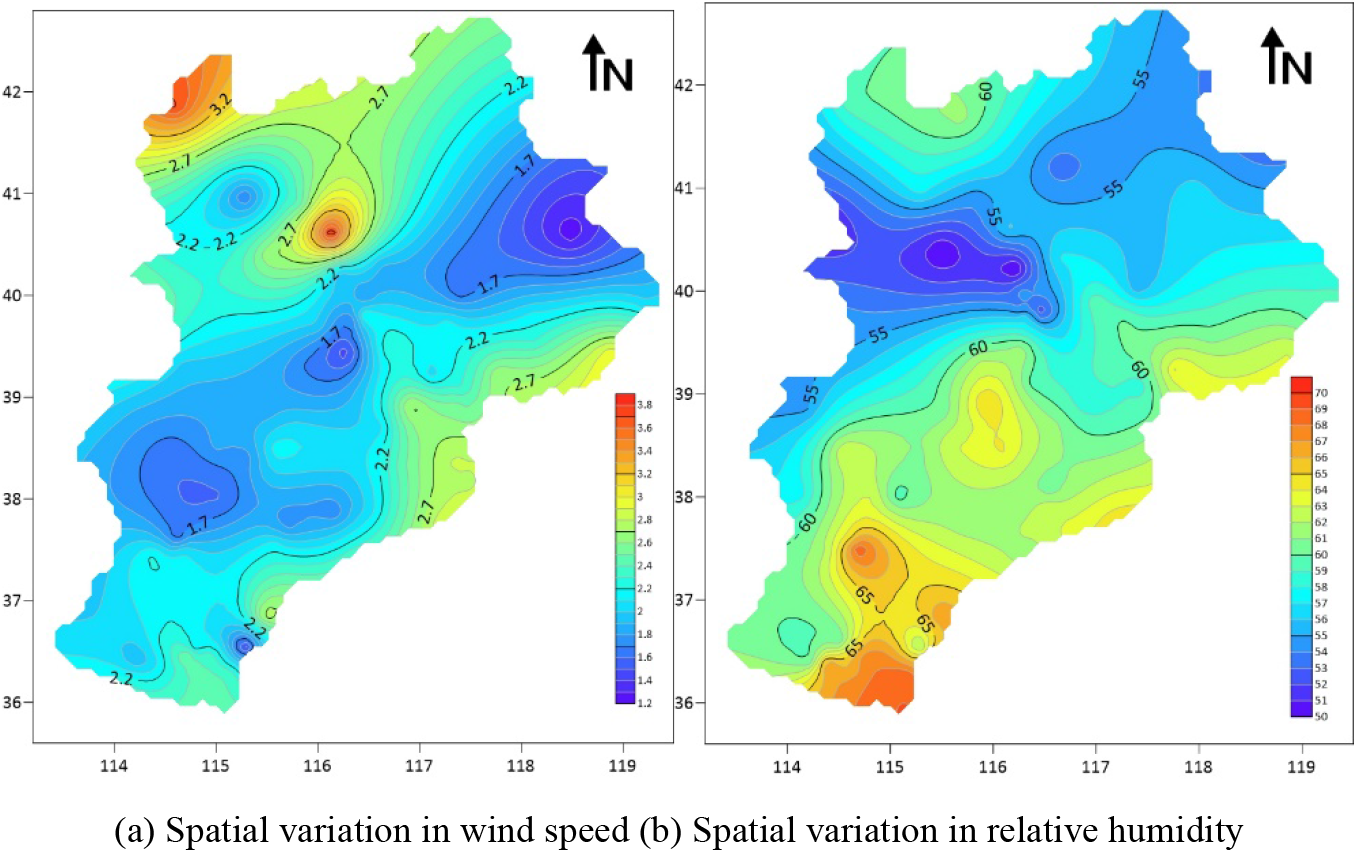
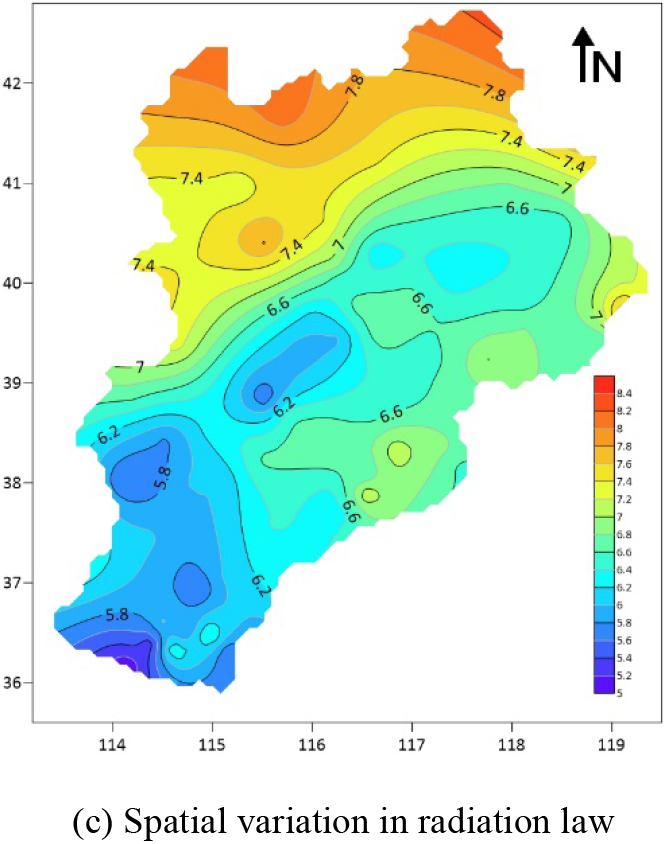
Map showing the characteristic analysis of the meteorological factors.

### 3.2 Distribution characteristics of potential evapotranspiration

Based on the maximum and minimum temperatures, relative humidity, solar radiation, wind speed, and other data from 80 meteorological stations in the Beijing-Tianjin-Hebei region, the potential evapotranspiration of the growth period was calculated using the FAO-98 procedure. The interannual variation of the potential evapotranspiration from 2000 to 2019 is shown in Figure 6. The plot of potential evapotranspiration in the region showed an overall upward trend that declined in certain years. For example, the potential evapotranspiration reached a minimum of 2.82.

**Figure 6.**
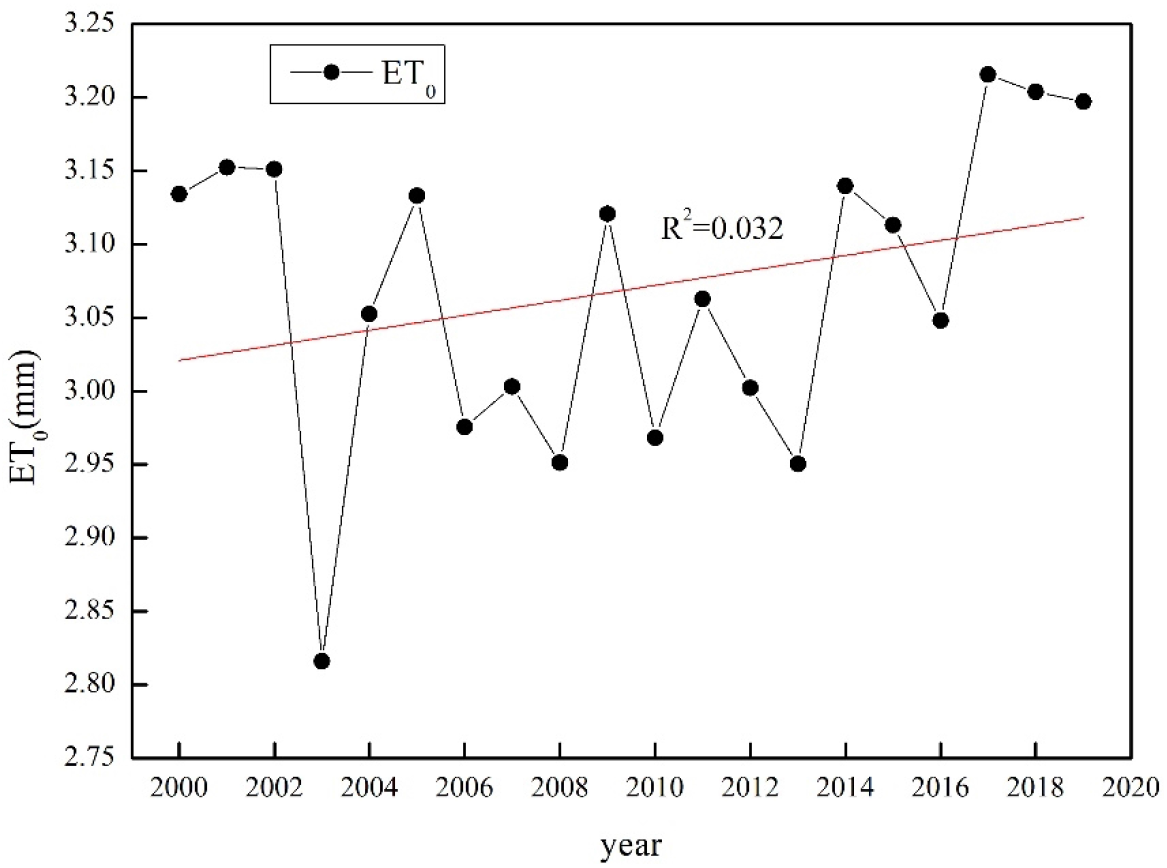
Plot showing the interannual variation of potential evapotranspiration in the growth period.

### 3.3 Distribution characteristics of effective precipitation

The interannual and spatial variations of effective precipitation in the Beijing-Tianjin-Hebei region from 2000 to 2019 were drawn, using the ten-year scale data of effective precipitation from 80 stations, as shown in Figure 7.

**Figure 7.**
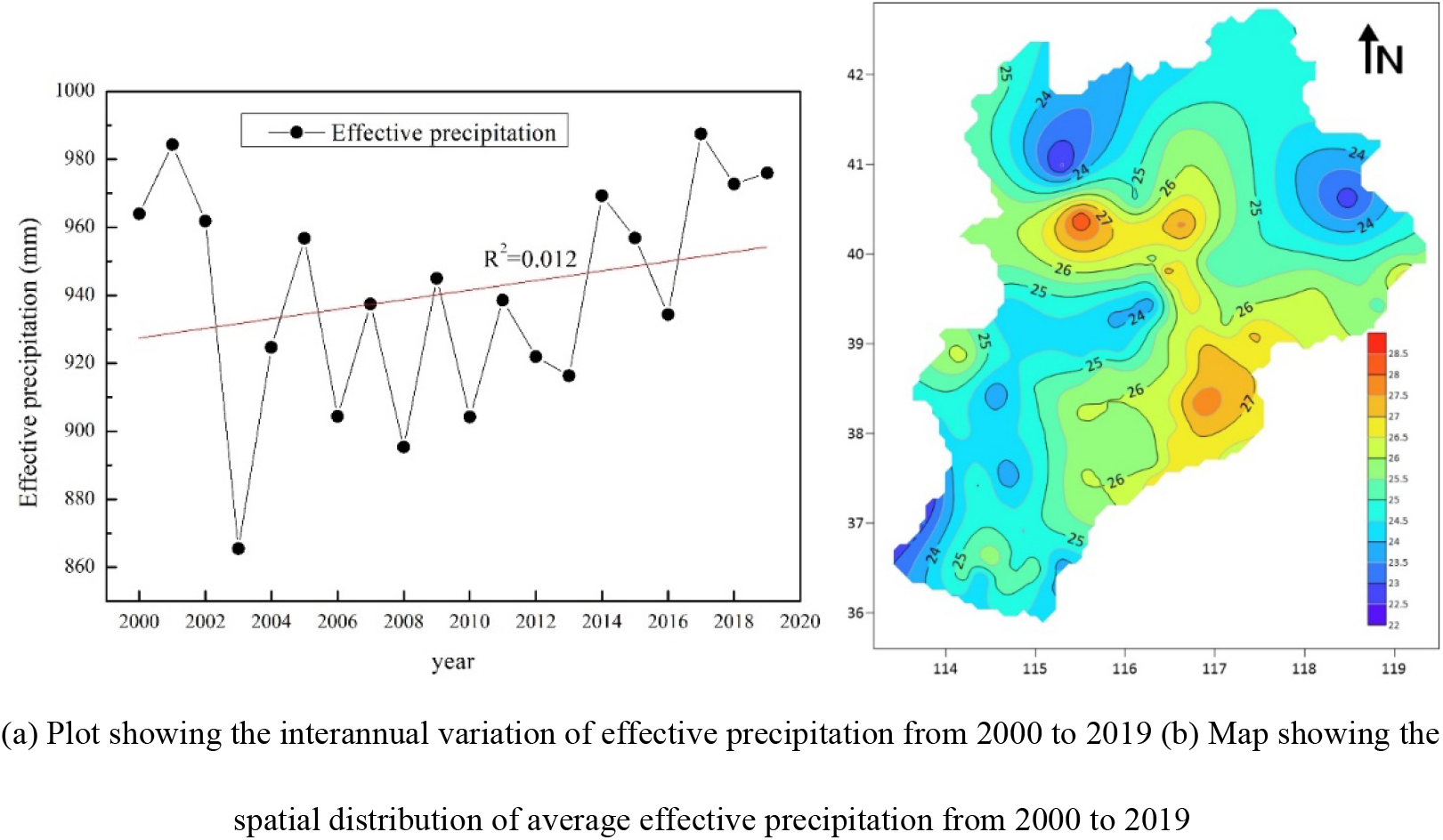
Distribution characteristics of effective precipitation in Beijing-Tianjin-Hebei region.

It can be concluded from the figure that the effective precipitation in the Beijing-Tianjin-Hebei region shows an upward trend. From the spatial distribution data of the 10-day average effective precipitation in the Beijing-Tianjin-Hebei region, obvious differences in the effective precipitation in the study area can be observed. In 2000, the effective precipitation was high in the eastern plains and relatively low in the northwest mountainous areas. In 2005, the effective precipitation at the boundary between mountains and plains was significantly low. By 2019, the Beijing-Tianjin-Hebei region had different years. The ten-day average effective precipitation was considerably high in the eastern plain.

### 3.4 Analysis of irrigation water requirement

#### 3.4.1 Analysis of the change characteristics

The changes in the irrigation water requirement of wheat at 80 sites from 2000 to 2019 were analyzed as shown in Figure 8(a). The overall trend of the plot showing irrigation water requirement of wheat in the study area is flat for most years. Fluctuations can be seen in 2002, 2006, and 2014, when the irrigation water requirement was relatively high or low. Figure 8(b) shows uneven spatial distribution of irrigation water requirement, that is high in the eastern plain and relatively low in the northwest mountainous area.

**Figure 8.**
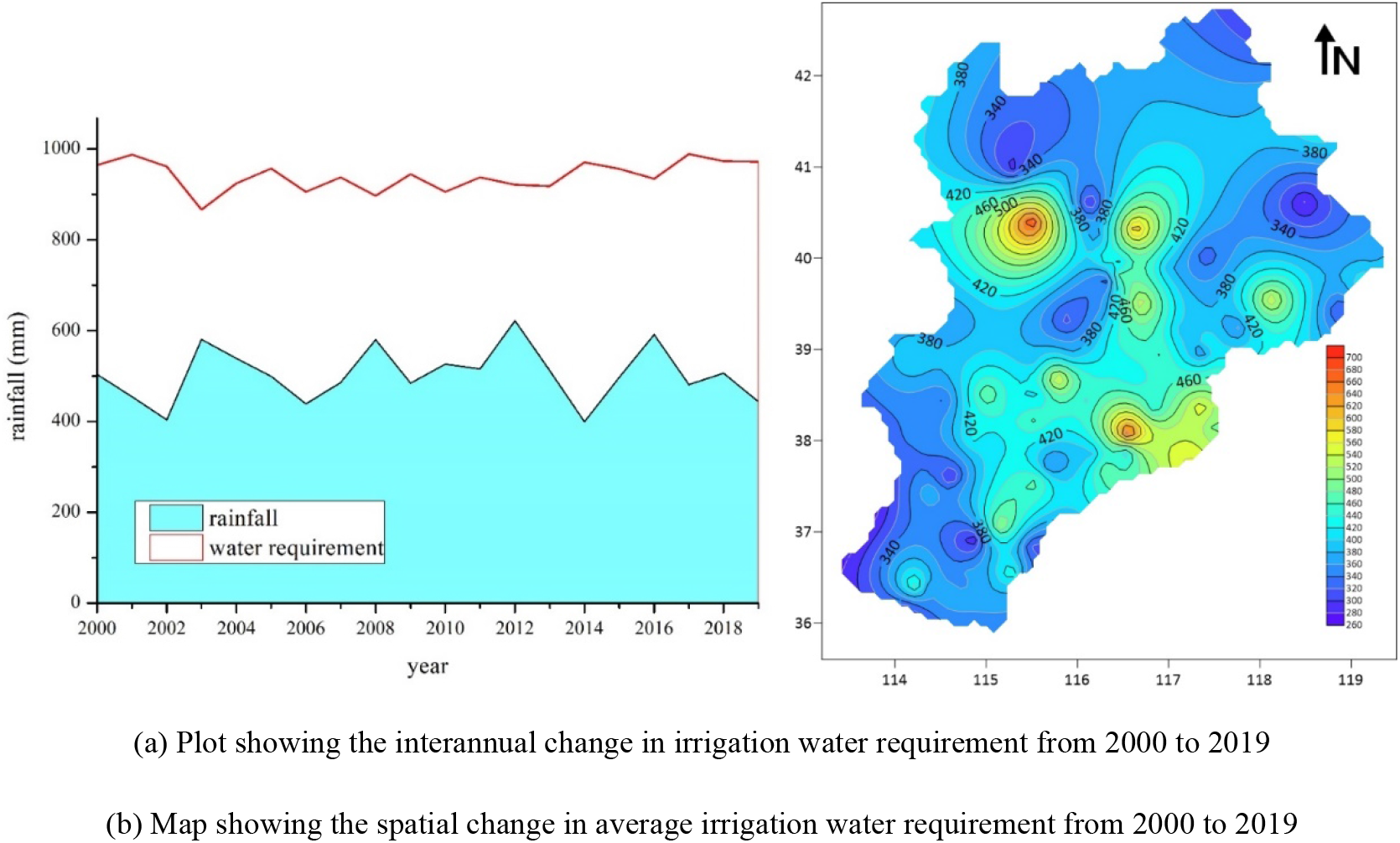
Distribution characteristics of irrigation water requirement in Beijing-Tianjin-Hebei region.

#### 3.4.2 Correlation analysis

The radar chart was used to analyze the changes in irrigation water requirement of wheat from 2000 to 2019, as shown in Figure 9. It can be clearly seen from the shape of the chart that irrigation water requirement in the study area is closest to potential evapotranspiration. However, for individual years, wind speed, relative humidity, and radiation contributions were different. The irrigation water requirement shows positively correlation with potential evapotranspiration, wind speed and radiation changes. The radar chart of the multi-year changes of different meteorological factors in the study area, showed that the overall change in irrigation water requirement has the highest correlation with potential evapotranspiration. However, for individual years, wind speed, relative humidity, and radiation were the common influencing factors affecting regional irrigation water requirement.

**Figure 9.**
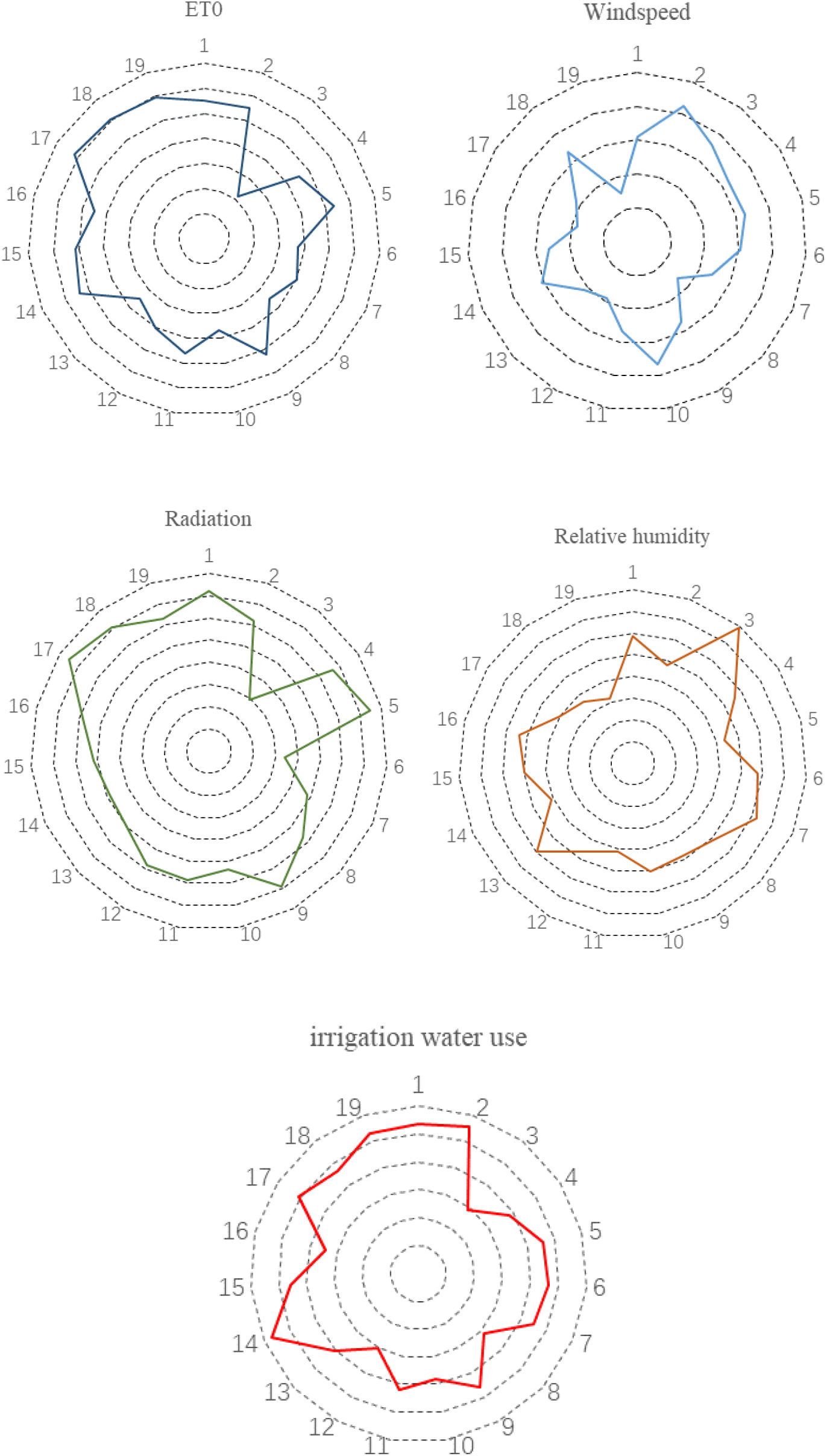
Radar chart showing changes in meteorological factors from 2000 to 2019.

## 4. Conclusion and discussion

### 4.1 Conclusion

(1) The annual average precipitation, average wind speed, and average relative humidity in the Beijing-Tianjin-Hebei region show a general downward trend, while the annual average temperature shows an upward trend. Relative humidity and radiation show negative spatial correlation, that is, areas with high relative humidity have low radiation, and vice versa.

(2) The irrigation water requirement of wheat in the Beijing-Tianjin-Hebei region gradually decreases spatially from north to south and from east to west. The temporal variation in irrigation water requirement in the eastern and western regions shows an opposite trend. The eastern region shows a gradually increasing trend, while the area is decreasing.

(3) Meteorological elements have significant impact on irrigation water requirement. They show negative correlation with irrigation water requirement, potential evapotranspiration, effective precipitation, and relative humidity, and positive correlation with sunshine hours, average temperature, and wind speed. The overall change in irrigation water requirement has the highest correlation with potential evapotranspiration. However, for yearly variations in regional irrigation water requirement, wind speed, relative humidity, and radiation are the common influencing factors.

### 4.2 Discussion

Irrigation water requirement is affected by complex factors. This study considers the meteorological factors influencing irrigation water requirement in different growth periods of wheat. Potential evapotranspiration and effective precipitation are observed to be the most important indicators that affect irrigation water requirement during the growth period. The effective precipitation during the wheat growth period in the Beijing-Tianjin-Hebei region has increased in the past two decades. This is contrary to the decreasing precipitation trend in the study area, as precipitation in the key growth periods of wheat has significantly increased since the 21st century. The study indicates that irrigation water requirement is affected by several factors at different growth stages of crops. A decrease in sunlight reduces the amount of energy reaching the surface of the Earth, resulting in a decrease in ground evaporation. Furthermore, a decrease in wind speed reduces the rate of water exchange between the air and the soil, affecting the soil moisture balance.

Most previous studies are based on the overall growth period of crops. However, this study focuses on each growth stage of the crop, in combination with meteorological elements and growth period data for comprehensive and accurate analyses. In future studies, these results of irrigation water requirement may be combined with groundwater extraction and artificial channel water transfer in the study area to increase crop yield.

Estimating the irrigation water requirement during the growth period of wheat is conducive to the adoption of targeted measures to effectively respond to climate change and increase wheat production. Water can thus be supplemented on time during the critical water requirement period, improving water use efficiency and crop yield.

## Availability of data

In this paper, we mainly used the stations data, wheat growth data data to support the findings of this study were supplied by the National Meteorological Information Center (http://data.cma.cn/) under license and so cannot be made freely available.

## Author contributions

Xu Chengcheng and Lu Chuiyu designed research; Lu Chuiyu performed research; Xu Chengcheng analyzed data; Wang Jianhhua contributed to interpretation of results; Lu Chuiyu contributed to algorithm development; and Xu Chengcheng wrote the paper.

## Competing interests

This manuscript has not been published or presented elsewhere in part or in entirety and is not under consideration by another journal. We have read and understood your journal’s policies, and we believe that neither the manuscript nor the study violates any of these. There are no conflicts of interest to declare.

## Acknowledgments

We acknowledge reviewers and editors for their patient and valuable advice on improving the quality of this paper and teaching us how to write higher quality papers. We thank our partners from Anhui University of Science and Technology and China Institute of Water Resources and Hydropower Research for their collaborative support during the studies. Financial support for this work was provided by the National Key Research and Development Program of China (grant No. 2016YFC0401404), Applied Technology Research and Development Program of Heilongjiang Province (grant No.GA19C005), National Key Research and Development Program (2016YFC0401300), and the National Science Fund for Distinguished Young Scholars (51625904).

